# Prider – multiplexed primer design using linearly scaling approximation of set coverage

**DOI:** 10.1101/2021.09.06.459073

**Authors:** Niina Smolander, Manu Tamminen

## Abstract

Designing oligonucleotide primers and probes is one of the key steps of various laboratory experiments such as multiplexed PCR or digital multiplexed ligation assays. When designing primers and probes to complex, heterogeneous DNA data sets, an optimization problem arises where the smallest number of oligonucleotides covering the largest diversity of the input dataset needs to be identified. Tools that provide this optimization in an efficient manner for large input data are currently lacking.

Here we present Prider, an R package for designing primers and probes with a nearly optimal coverage for complex and large sequence sets. Prider initially prepares a full primer coverage of the input sequences, the complexity of which is subsequently reduced by removing components of high redundancy or narrow coverage. The primers from the resulting near-optimal coverage are easily accessible as data frames and their coverage across the input sequences can be visualized as heatmaps using Prider’s plotting function. Prider scales linearly to increasing sequence data and therefore permits efficient design of primers to large and highly diverse DNA datasets.

Prider is available on GitHub under the permissive BSD-3-clause license:

https://github.com/manutamminen/prider

## Introduction

Multiplex molecular techniques, such as multiplex polymerase chain reaction (PCR) (Chamberlain et al., 1988) and digital multiplex ligation assay (dMLA) (Tamminen et al., 2020), are methods for detecting multiple genomic targets in a single experiment. These techniques have enabled the development of various screening methods in the fields such as pathogen detection and human genetics and utilize sets of primers or probes that can detect even hundreds of targets (Teder *et al*., 2018; Kosztolányi *et al*., 2018; Benard-Slagter *et al*., 2017; Grigorenko *et al*., 2014; Makowski *et al*., 2003).

Designing primers or probes for optimal detection of multiple targets in complex and large sets of DNA sequences is an instance of set coverage problems which aim to find a minimal set of primer sequences that cover the total DNA sequence set (Shyu and Lee, 1990). Various tools have been created for multiplex primer and probe designing, such as the command line based *PriMux* (Hysom *et al*., 2012), the web-application *PrimerDesign* (Brodin *et al*., 2013), the R package DECIPHER’s *DesignPrimers* and *DesignProbes* (Wright *et al*., 2014), the GUI *PrimerMapper* (O’Halloran, 2016) and the R package *openPrimeR* (Kreer *et al*., 2020). However, these tools are either poorly suited for scripting and/or scale poorly to large input sequence sets.

Here we present an R package Prider, which prepares a nearly optimal primer coverage for an input FASTA file and scales linearly to increasing sequence data. Prider is a flexible tool that can be used for both designing primers and probes for highly scalable molecular screening and quantification applications (Tamminen et al., 2020; Teder *et al*., 2018; Kosztolányi *et al*., 2018; Benard-Slagter *et al*., 2017;). It gives the user an ability to control variety of primer/probe properties and is well suited for scripting. The key features of Prider are its linear scalability to large and complex datasets, capability of approximating near-optimal set coverage with minimal user intervention, as well as its inbuilt ability to visualize the output.

## Implementation

### Input and parameters

Prider was developed on R version 4.0.5 (R Core Team, 2021) with the package *Rcpp* 1.0.7 (Eddelbuettel and François, 2011) using C++11. The input to Prider is a single FASTA file, containing the sequences to which primers or probes are to be designed. User can change the primer length, the minimum primer and sequence group sizes and the number of cumulative coverage decimals, explained below. Furthermore, additional filtering via GCcheck parameter removes the primers with proportional G and C base contents outside the range set by GCmin and GCmax parameters. Filtering via GChalves parameter removes the primers that have too large a difference in proportional GC contents between the two halves of the primer. The maximum difference is set by GCsimilarity parameters.

### Cluster preparation and filtering

The first step of primer cluster preparation is the division of each DNA sequence from the input FASTA file into sub-sequences - primer candidates - using a sliding window function *chunker* implemented in C++. During the process the primer candidates remain associated with their respective FASTA headers. Primer candidates that include any other characters than A, G, C or T are removed. Subsequently, primer candidates shared by multiple input sequences are used to group together sequences with shared motifs. These sequence groups are further grouped together, linking different primer candidates together and producing a data frame of all sequence clusters and primer clusters which cover them.

To optimize the number of primer candidates needed to cover the input FASTA, the primer clusters with sequence coverage or sequence cluster size below user-defined cut-off are excluded. The primer clusters are subsequently ordered by their size and the cumulative contribution of each cluster to the total sequence coverage are calculated. Those clusters whose cumulative contribution is below user-defined cut-off are excluded. Primer clusters with a similar cumulative coverage are grouped together and only the primer cluster with the largest sequence and primer group sizes within each cumulative coverage group is kept. The resulting data frame contains a character vector of the primer groups providing the near-optimal sequence coverage, and a respective character vector of the covered input sequences.

### Prider output

The output of Prider is an S3-decorated list with five elements accessible with either Prider’s S3 methods, indexing, or the “$” operator. The output elements are:

1. Description; summarizes the contents of the input FASTA and the produced Prider list.
2. Conversion table; a data frame that contains the original FASTA headers, full DNA sequences and the sequence ids.
3. Primer candidates; a data frame that contains the primer group DNA sequences, an identification number for each primer group, the sequence ids associated with the primer clusters, primer cluster and sequence group sizes and the cumulative coverage values.
4. Excluded sequences; a data frame with the same format as conversion table. Contains the sequences, which were not associated with any primer cluster due to filtering criteria.
5. Primer matrix; a TRUE-FALSE table where each row is a primer group and each column is a single sequence id. This is the input for the S3 plotting function for the Prider objects.

## Results and conclusion

Two randomly generated FASTA test sets were used to evaluate the processing speed of Prider; one with increasing total number of bases per file (300 sequences each) and one with increasing number of sequences per file (465000 bases each). The sets consisted of 310 and 300 files, respectively, and 10 replicates of each number of bases or sequences. To make sure that even the smallest files could be processed, the parameter *minimum_sequence_group_size* was set to 1.

The processing time of Prider (determined by the *user*.*self* value of the base R function *system*.*time*) was linearly dependent on the number of input bases, with 3e4 bases distributed to 300 sequences taking approx. 0.5 seconds and 9.03e6 bases taking approx. 300 seconds (fig. 1A) on a Macbook Pro (M1, 8 GB, 2020, macOS Big Sur). The number of sequences the bases were distributed on had a minor, irregular effect on the processing time (fig. 1B). The test data and the code used for the tests is available at https://zenodo.org/record/5361309. The benchmarks reveal that Prider scales well to large sequence data and has low variation between the processing times of the replicates.

**Figure 1.**
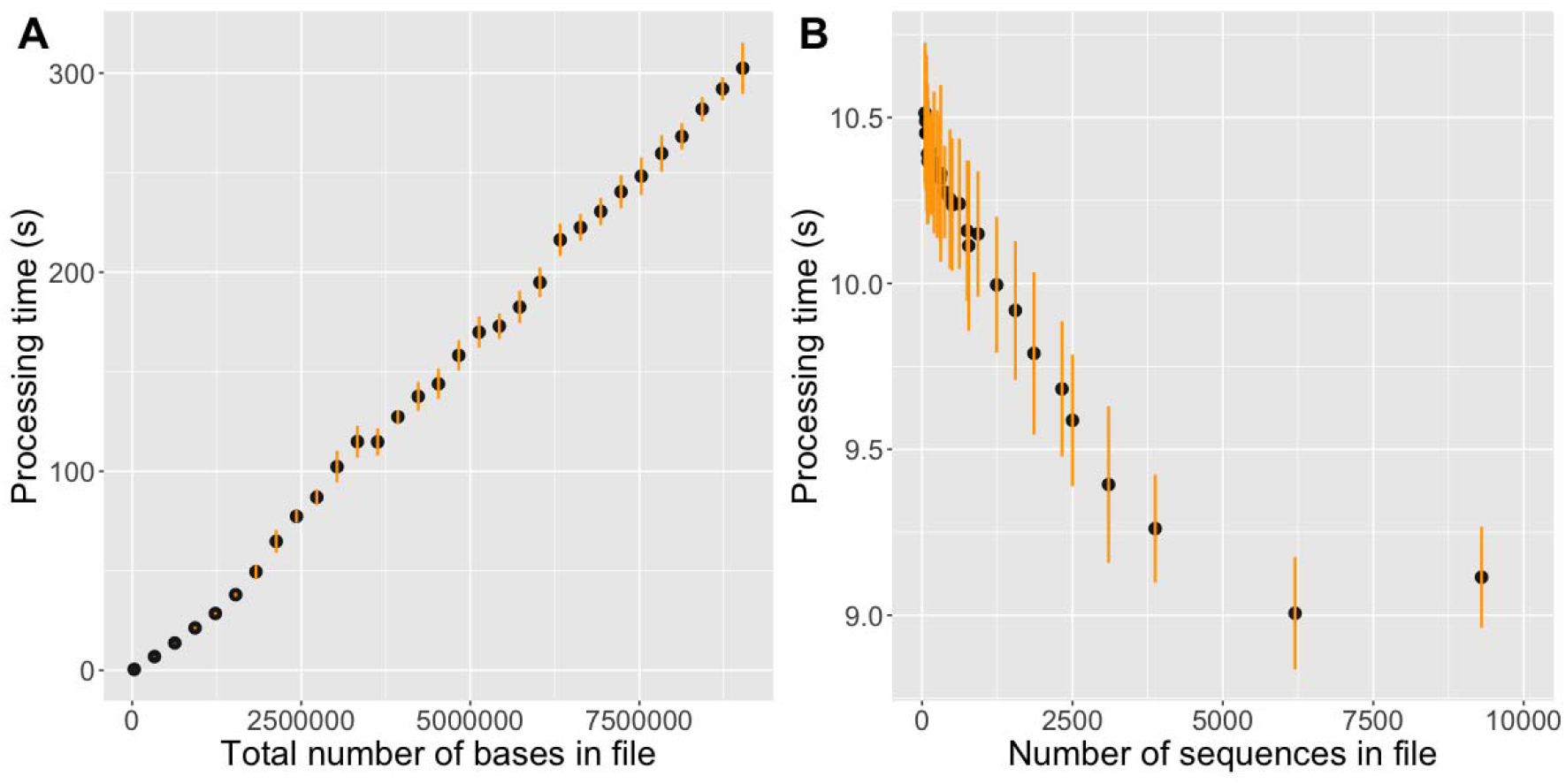
Prider processing time (dot) and standard deviation (line) for A) varying numbers of bases in files, B) varying numbers of sequences in files (total 465000 bases each).

Design of multiplexed primers and probes to highly diverse DNA data is a problem commonly encountered in various screening applications (Tamminen *et al*, 2020; Kosztolányi *et al*., 2018; Teder *et al*., 2018; Benard-Slagter *et al*., 2017). For instance, in antimicrobial resistance surveillance one needs to account for the extremely high diversity of antibiotic resistance genes which have been reported in various environments (Brandt *et al*., 2017). A similar situation arises in pathogenicity detection where the reported diversity of known pathogenicity genes is high (Yoon et al., 2015). Such screening applications greatly benefit from Prider since its linear scalability permits processing sequence data of this scale. Thus, combination of Prider with highly scalable molecular quantification techniques such as dMLA (Tamminen *et al*, 2020) will permit an unprecedented molecular screening capability with immediate applicability in fields such as clinical microbiology or antimicrobial resistance surveillance.

## Acknowledgements

The authors acknowledge Dr. Tim Julian (Eawag, Switzerland) for ideas which were instrumental in shaping the described method.

